# The clinical and functional effects of *TERT* variants in myelodysplastic syndrome

**DOI:** 10.1101/2021.02.11.430624

**Authors:** Christopher R. Reilly, Mikko Myllymäki, Robert Redd, Shilpa Padmanaban, Druha Karunakaran, Valerie Tesmer, Frederick D. Tsai, Christopher J. Gibson, Huma Q. Rana, Liang Zhong, Wael Saber, Stephen R. Spellman, Zhen-Huan Hu, Esther H. Orr, Maxine M. Chen, Immaculata De Vivo, Corey Cutler, Joseph H. Antin, Donna Neuberg, Judy E. Garber, Jayakrishnan Nandakumar, Suneet Agarwal, R. Coleman Lindsley

## Abstract

Germline pathogenic *TERT* variants are associated with short telomeres and an increased risk of developing myelodysplastic syndrome (MDS) among patients with a telomere biology disorder. We identified *TERT* rare variants in 41 of 1514 MDS patients (2.7%) without a clinical diagnosis of telomere biology disorder who underwent allogeneic transplantation. Patients with *TERT* rare variants had shorter telomere length (p<0.001) and younger age at MDS diagnosis (52 vs. 59 years, p=0.03) than patients without a *TERT* rare variant. In multivariable analyses, *TERT* rare variants were associated with inferior overall survival (p=0.034) driven by an increased incidence of non-relapse mortality (NRM) (p=0.015). Death from a non-infectious pulmonary cause was more frequent among patients with a *TERT* rare variant. According to ACMG/AMP guidelines and Sherloc criteria, 39 *TERT* rare variants were classified as VUS and one as likely pathogenic. Therefore, we cloned all rare missense variants and quantified their impact on telomere elongation in a cell-based assay. We found that 36 of 40 variants had severe or intermediate impairment in their capacity to elongate telomeres. Using a homology model of human TERT bound to the shelterin protein TPP1, we inferred that TERT rare variants disrupt domain-specific functions, including catalysis, protein-RNA interactions, and recruitment to telomeres. Our results indicate that the contribution of *TERT* rare variants to MDS pathogenesis and NRM risk is underrecognized and routine screening for *TERT* rare variants in MDS patients regardless of age or clinical suspicion could identify clinically inapparent telomere biology disorders and improve transplant outcomes through risk-adapted approaches.

## INTRODUCTION

Impaired telomere maintenance is implicated in the pathogenesis of myelodysplastic syndrome (MDS),^1–5^ for which allogeneic hematopoietic stem cell transplantation (HSCT) is the only potential cure.^6,7^ Shorter pre-transplant blood telomere length is independently associated with an increased risk of early non-relapse mortality (NRM) in MDS patients,^8^ but the genetic determinants of telomere length in MDS are incompletely characterized.

Germline pathogenic variants affecting telomerase- and telomere-associated proteins cause a global impairment in telomere maintenance and short telomeres in all tissues.^1,9–11^ Individuals with dyskeratosis congenita (DC), an early-onset syndromic presentation of a telomere biology disorder, have characteristic mucocutaneous features, bone marrow failure,^2,12^ and a markedly increased risk of developing MDS or acute myeloid leukemia.^2,4,13,14^ In contrast, adult patients with a telomere biology disorder more frequently present with aplastic anemia,^15–17^ idiopathic pulmonary fibrosis,^18–23^ and liver cirrhosis.^21,24,25^ Affected families may show anticipation, marked by changes in the onset, phenotype, and severity of clinical disease across successive generations.^21,26–28^ Clinical suspicion for telomere biology disorder is based on the presence of syndromic features and disease phenotypes in relatives.^3,22^ However, clinical manifestations are highly variable, and up to 40% of affected patients lack a family history of hematologic, pulmonary, or hepatic abnormalities.^4^ In contrast to DC, the risk of developing MDS in older adults with late-presenting or unrecognized telomere biology disorders is unknown.

*TERT* is the most frequently mutated gene among patients with a telomere biology disorder^4^ and can cause disease in an autosomal dominant form.^1^ *TERT* encodes telomerase reverse transcriptase that binds to the telomerase RNA component (TERC) and functions within the multi-subunit telomerase holoenzyme complex to extend telomere ends during DNA replication.^9,29^ Telomerase is composed of four structural domains with distinct functional roles:^30,31^ the telomerase essential/N-terminal (TEN) domain, telomerase RNA-binding domain (TRBD), reverse transcriptase domain (RTD), and C-terminal extension (CTE) domain. Disease-associated germline *TERT* variants are predominantly missense substitutions and occur within all structural domains.^32^ Novel *TERT* missense variants are classified as variants of unknown significance (VUS) by consensus guidelines in the absence of additional supporting computational or functional evidence of pathogenicity.^33,34^ However, *in silico* prediction algorithms have limited utility in assessing genotype-phenotype relationships for missense substitutions.^35–37^ Furthermore, the cellular effect of *TERT* variants may not be reflected by *in vitro* functional assays.^38–40^

The prevalence and clinical significance of *TERT* variants among MDS patients unselected for suspicion of a telomere biology disorder are unknown. Here we analyzed the clinical and functional effects of *TERT* variants in a registry-level cohort of MDS patients who underwent allogeneic HSCT.

## RESULTS

### TERT variants in MDS and NHL cohorts

In total, we identified 270 nonsynonymous *TERT* coding variants among the MDS and NHL cohorts (Figure 1A). Pathogenic genetic variants are observed infrequently in the general population due to strong negative selection.^34,49^ Therefore, we grouped *TERT* variants based on their maximum gnomAD population allele frequency in any reference population,^50^ where “common” variants had a maximum allele frequency ≥0.001 and “rare” variants had a maximum allele frequency <0.001. Using this approach, 228 variants (84.4%) were classified as common and 42 variants (R1086H occurred in two MDS patients) as rare (15.6%) (Figure 1A). The frequency of *TERT* common variants was similar in the MDS and NHL cohorts (11.9% vs. 12.0%, p=0.79), and primarily included the SNPs p.A279T (rs61748181), p.H412Y (rs34094720), p.E441del (rs377639087), and p.A1062T (rs35719940) (Figure 1C and Figure S1). In contrast, *TERT* rare variants were significantly more common in patients with MDS (2.7% MDS vs. 0.25% NHL, p <0.001)(Figure 1B).

**Figure 1.**
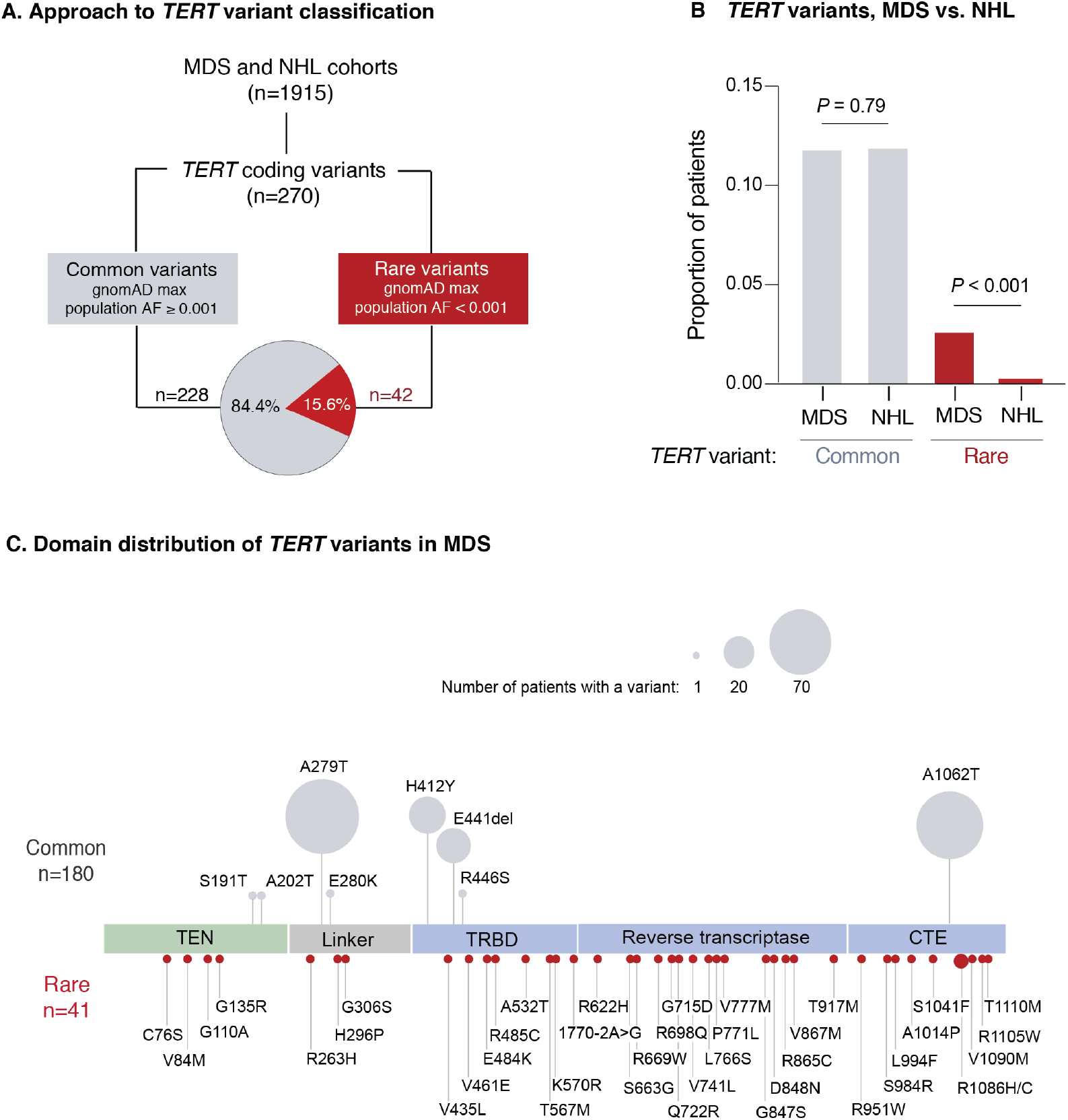
*TERT* variants in MDS and NHL. A) Classification approach of nonsynonymous *TERT* variants identified in the MDS and NHL cohorts. B) Frequency of *TERT* common and *TERT* rare variants within the MDS and NHL cohorts. C) Domain distribution of *TERT* variants within the MDS cohort. *TERT* common variants (n= 180 variants) and rare variants (n=40 variants among 41 patients) are located above and below the coding region, respectively. The size of each ball is proportional to the number of patients with that variant. *TERT* rare variants are colored in red and *TERT* common variants in gray.

In MDS patients, *TERT* rare variants occurred within all structural domains: RTD (n=15), CTE domain (n=10), TRBD (n=8), TEN domain (n=4), and linker region between TEN and TRBD (n=3) (Figure 1C). The majority of variants were missense substitutions (39 of 40) with one splice site variant (c.1770-2A>G). According to ACMG/AMP guidelines^33^ and Sherloc criteria^34^, 39 variants were classified as VUS and only one variant (p.R865C) was classified as likely pathogenic (Table 1). Twenty-three variants (57.5%) were absent from gnomAD, 17 variants were listed in ClinVar (42.5%), and 11 variants (28.2%) have been published in patients with telomere biology diseases. *In silico* Combined Annotation-Dependent Depletion (CADD)^49^ PHRED-like scores > 20 and ClinPred^51^ scores >0.5 were observed in 54% and 51% of variants, respectively (Table 1 and S2). In contrast, *TERT* common variants were classified as benign or likely benign (Table S3), and most have been experimentally determined to have comparable activity to wild type TERT.^40^

**Table 1.** *TERT* rare variant characteristics.

| *TERT* rare variant | Structural domain | gnomAD max popAF | ACMG/AMP Classification | Sherloc Classification | CADD PHRED score | ClinVar Accession Number |
| --- | --- | --- | --- | --- | --- | --- |
| p.C76S | TEN | 0 | VUS | VUS | 10.59 |  |
| p.V84M | TEN | 0 | VUS | VUS | 22.8 |  |
| p.G110A | TEN | 0 | VUS | VUS | 15.96 |  |
| p.G135R | TEN | 0.0008400 | VUS | VUS | 16.84 | VCV000410665 |
| p.R263H | Linker | 0 | VUS | VUS | 3.266 |  |
| p.H296P | Linker | 0.0002000 | VUS | VUS | 4.311 | VCV000268080 |
| p.G306S | Linker | 0 | VUS | VUS | 4.83 |  |
| p.V435L | TRBD | 0 | VUS | VUS | 6.863 |  |
| p.V461E | TRBD | 0 | VUS | VUS | 26.5 |  |
| p.E484K | TRBD | 0 | VUS | VUS | 2.57 | VCV000955018 |
| p.R485C | TRBD | 0.0000638 | VUS | VUS | 18.03 |  |
| p.A532T | TRBD | 0.0000265 | VUS | VUS | 10.15 | VCV000581635 |
| p.T567M | TRBD | 0 | VUS | VUS | 17.36 |  |
| p.K570R | TRBD | 0 | VUS | VUS | 23.7 |  |
| c.1770-2A>G | TRBD | 0 | VUS | VUS | 26.2 |  |
| p.R622H | RTD | 0 | VUS | VUS | 24.9 |  |
| p.S663G | RTD | 0 | VUS | VUS | 13.49 |  |
| p.R669W | RTD | 0.0000531 | VUS | VUS | 22.5 | VCV000539196 |
| p.R698Q | RTD | 0 | VUS | VUS | 23.5 |  |
| p.G715D | RTD | 0 | VUS | VUS | 24.3 |  |
| p.Q722R | RTD | 0 | VUS | VUS | 23.7 | VCV000471853 |
| p.V741L | RTD | 0.0002000 | VUS | VUS | 14.03 | VCV000652891 |
| p.L766S | RTD | 0 | VUS | VUS | 22.4 |  |
| p.P771L | RTD | 0 | VUS | VUS | 24.1 |  |
| p.V777M | RTD | 0 | VUS | VUS | 19.9 | VCV000436985 |
| p.G847S | RTD | 0.00006482 | VUS | VUS | 24.4 |  |
| p.D848N | RTD | 0.0000089 | VUS | VUS | 22.8 |  |
| p.R865C | RTD | 0.0000240 | LP | LP | 24.6 | VCV000986922 |
| p.V867M | RTD | 0.0000000 | VUS | VUS | 22.6 | VCV000242683 |
| p.T917M | RTD | 0.0001240 | VUS | VUS | 21.5 | VCV000857994 |
| p.R951W | CTE | 0.000008833 | VUS | VUS | 20.9 | VCV000836202 |
| p.S984R | CTE | 0.000008828 | VUS | VUS | 15.71 |  |
| p.L994F | CTE | 0.00002716 | VUS | VUS | 22.7 | VCV000580043 |
| p.A1014P | CTE | 0 | VUS | VUS | 24.5 |  |
| p.S1041F | CTE | 0 | VUS | VUS | 17.63 |  |
| p.R1086C | CTE | 0 | VUS | VUS | 15.92 |  |
| p.R1086H | CTE | 0.000839 | VUS | VUS | 20.6 | VCV000242237 |
| p.V1090M | CTE | 0.0003000 | VUS | VUS | 15.67 | VCV000012733 |
| p.R1105W | CTE | 0 | VUS | VUS | 22.2 | VCV000939229 |
| p.T1110M | CTE | 0.0002000 | VUS | VUS | 12.05 | VCV000039122 |

### TERT rare variants and clinical characteristics

Patients with a *TERT* rare variant had shorter telomere length (ddCT=0.405 vs. 0.507, p<0.001) and were younger at MDS diagnosis (median age 52 vs. 59, p=0.03) than those without a *TERT* variant (Figure 2A and B). All other clinical characteristics, including Karnofsky performance status, hematopoietic stem cell transplant comorbidity index (HCT-CI) score, peripheral blood counts at transplant, frequency of therapy-related MDS, and IPSS-R risk group were similar in patients with and without a *TERT* rare variant (Table 2 and Table S6). Patients with a *TERT* common variant were similar to those without any *TERT* variant with respect to telomere length (ddCT=0.509 vs. 0.507, p=0.80), age at MDS diagnosis (median age 59 vs. 59, p=0.72), and other clinical characteristics (Figure 2A and B, Table S6).

**Table 2.** Patient characteristics by *TERT* rare variant status.

|  | No *TERT* rare<br>n = 1473 | *TERT* rare<br>n = 41 | p-value |
| --- | --- | --- | --- |
| **Patient-related variables** |  |  |  |
| Age at transplantation, median (range), years | 59 (0 - 77) | 52 (14 - 72) | 0.03<sup>^</sup> |
| Female sex, n (%) | 591 (40) | 11 (27) | 0.11<sup>†</sup> |
| Karnofsky performance status score < 90, n (%) | 403 (27) | 16 (39) | 0.15<sup>†</sup> |
| Hematopoietic cell transplant comorbidity index |  |  | 0.15<sup>‡</sup> |
| 0 | 255 (25) | 3 (11) |  |
| 1-2 | 247 (24) | 8 (29) |  |
| 3 | 535 (52) | 17 (61) |  |
| Missing | 436 | 13 |  |
| **Disease-related variables** |  |  |  |
| Hemoglobin, median (IQR), g/dL | 9.4 (8.1-11.2) | 9.9 (8.6-11.1) | 0.26<sup>^</sup> |
| Platelet count, median (IQR), x 10<sup>9</sup>/L | 72 (30-147) | 72 (37-115) | 0.87<sup>^</sup> |
| Absolute neutrophil count, median (IQR), x 10<sup>9</sup>/L | 1.1 (0.5-2.3) | 1.3 (0.5-2.6) | 0.63<sup>^</sup> |
| Bone marrow blasts at transplant, median (IQR), % | 3 (1-6) | 1 (0-5) | 0.03<sup>^</sup> |
| Prior MDS-directed therapy, n (%) | 861 (58) | 24 (59) | 0.99<sup>†</sup> |
| Therapy-related MDS, n (%) | 305 (21) | 6 (15) | 0.43<sup>†</sup> |
| **Somatic mutations** |  |  |  |
| *ASXL1* | 289 (20) | 8 (20) | 0.99<sup>†</sup> |
| *U2AF1* | 119 (8) | 8 (20) | 0.02<sup>†</sup> |
| *TP53* | 282 (19) | 7 (17) | 0.84<sup>†</sup> |
| *ETV6* | 57 (4) | 5 (12) | 0.02<sup>†</sup> |
| *RUNX1* | 169 (11) | 5 (12) | 0.81<sup>†</sup> |
| *PPM1D* | 84 (6) | 4 (10) | 0.29<sup>†</sup> |
| **Transplant-related variables** |  |  |  |
| Conditioning regimen, n (%) |  |  | 0.10<sup>†</sup> |
| Myeloablative | 765 (52) | 24 (59) |  |
| Reduced intensity | 565 (39) | 17 (41) |  |
| Nonmyeloablative | 130 (9) | 0 (0) |  |
| Missing | 13 | 0 |  |
| Donor type, n (%) |  |  | 0.85<sup>†</sup> |
| Matched, Related | 176 (12) | 5 (12) |  |
| Matched, Unrelated | 837 (57) | 26 (63) |  |
| Mismatched | 289 (20) | 7 (17) |  |
| Cord Blood | 171 (12) | 3 (7) |  |
| Graft type, n (%) |  |  | 0.88<sup>†</sup> |
| Bone marrow | 215 (15) | 6 (15) |  |
| Peripheral blood stem cells | 1,082 (73) | 32 (78) |  |
| Cord Blood | 165 (11) | 3 (7) |  |
| Other | 11 (1) | 0 (0) |  |

<sup>^</sup>Wilcoxon rank-sum test,<sup>†</sup>Fisher's exact test,<sup>‡</sup>Cochran-Armitage trend test
Peripheral blood counts and bone marrow blast counts at time of transplantation
IQR, interquartile range

**Figure 2.**
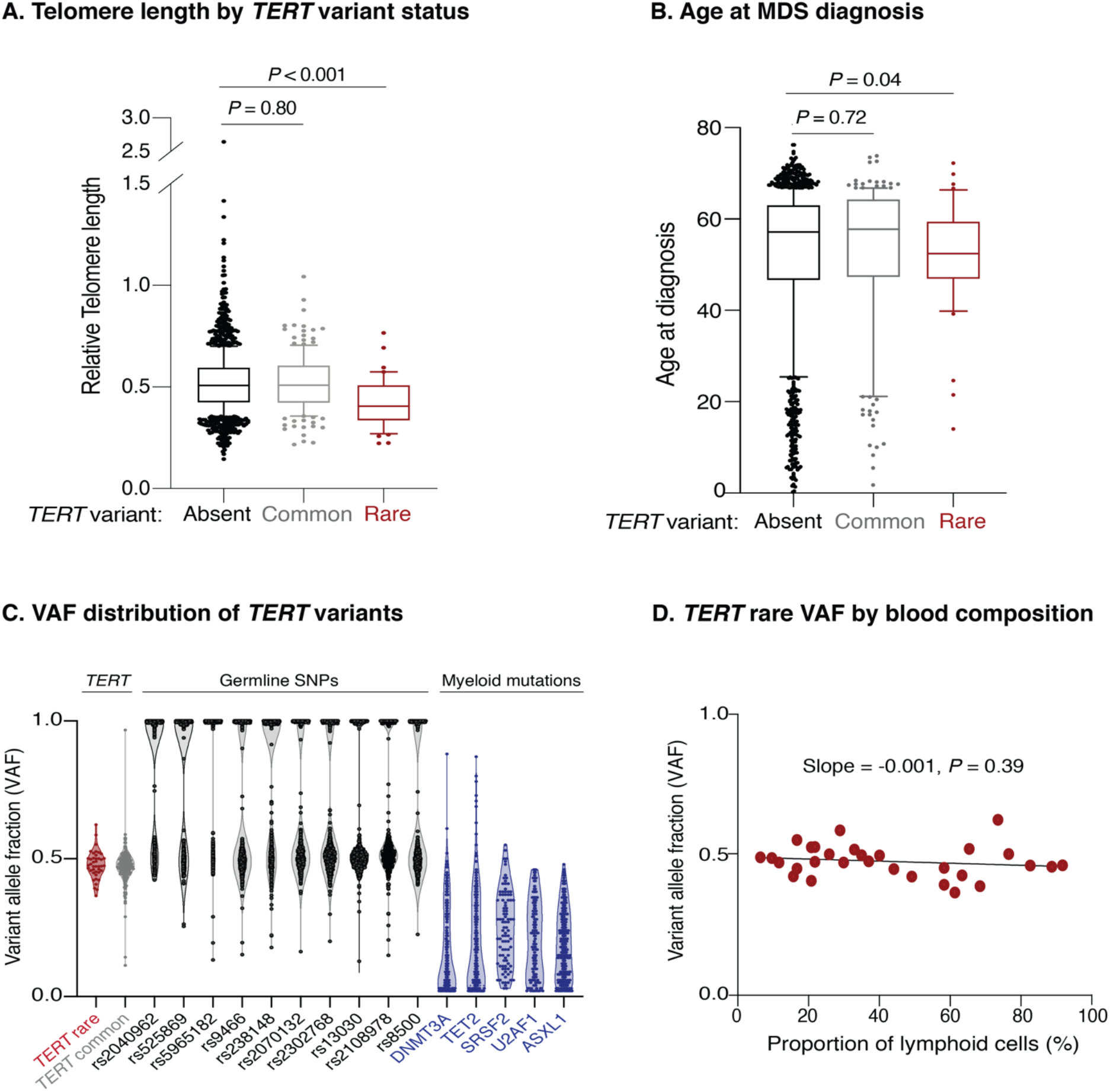
Association of TERT variants with telomere length and age at MDS diagnosis. Pre-transplant whole blood relative telomere length by *TERT* variant status. B) Age at MDS diagnosis by *TERT* variant status. *TERT* variant groups are labelled as follows: no *TERT* variant (black), *TERT* common variant (gray), and *TERT* rare variant (red). C) Variant allele fraction distribution of *TERT* variants, germline SNPs (black), and myeloid mutations (blue). D) Variant allele fraction distribution of *TERT* rare variants as a function of the proportion of blood lymphocytes. Linear regression slope with p-value is shown.

In the absence of available constitutional reference tissue, we used genetic characteristics to determine whether *TERT* rare variants were likely present in the germline or acquired somatically within the malignant clone. Germline variants are present in all cells and have a variant allele fraction (VAF) around 0.5 (heterozygous) or 1 (homozygous), while somatic mutations are present only in a subset of clonal cells with a wider range of VAF that falls below 0.5 in diploid cells. *TERT* rare variants and control germline SNPs had a median VAF of 0.48 (95% CI 0.457-0.502) and 0.56 (95% CI 0.557-0.564), respectively (Figure 2C), whereas MDS somatic mutations typically present in the founding clone displayed lower median VAFs (*DNMT3A*: 0.10, *TET2*: 0.16, *SRSF2*: 0.25, *U2AF1*: 0.20, *ASXL1*: 0.18) Additionally, the VAF of *TERT* rare variants did not vary with the proportion of blood lymphocytes (Figure 2D), indicating that the variants were present in all nucleated cells, including both the clonal myeloid compartment and the typically non-clonal lymphoid compartment.

### TERT rare variants and clinical outcomes

To determine whether *TERT* variant status was associated with clinical outcomes after transplantation, we evaluated overall survival and the cumulative incidences of relapse and NRM. Among 41 MDS patients with a *TERT* rare variant, overall survival (OS) was 24% at 5 years (Figure 3A), and the cumulative incidences of NRM and relapse at 5 years were 52.5% and 27.5%, respectively (Figures 3B and C). In multivariable analysis, the presence of a *TERT* rare variant compared with the absence of a *TERT* rare variant was associated with inferior overall survival (HR for death 1.50, 95% CI 1.04-2.20, p=0.03) and an increased rate of NRM (HR 1.75, 95% CI 1.13-2.72, p=0.01) but not a higher rate of relapse (HR 0.78, 95% CI 0.42-1.16, p=0.44) (Figure 3D). Non-genetic factors also impacted the rate of non-relapse mortality in this model, including recipient age (per 10 year increase, HR 1.23, 95% CI 1.14-1.33, p<0.01) and Karnofsky performance score <90 (1.23, 1.02-1.53, p=0.03). The results of the multivariable Cox model for overall survival and the Fine–Gray model for the rates of relapse and NRM, along with adjusted covariates for each model, are provided in Tables S8.

**Figure 3.**
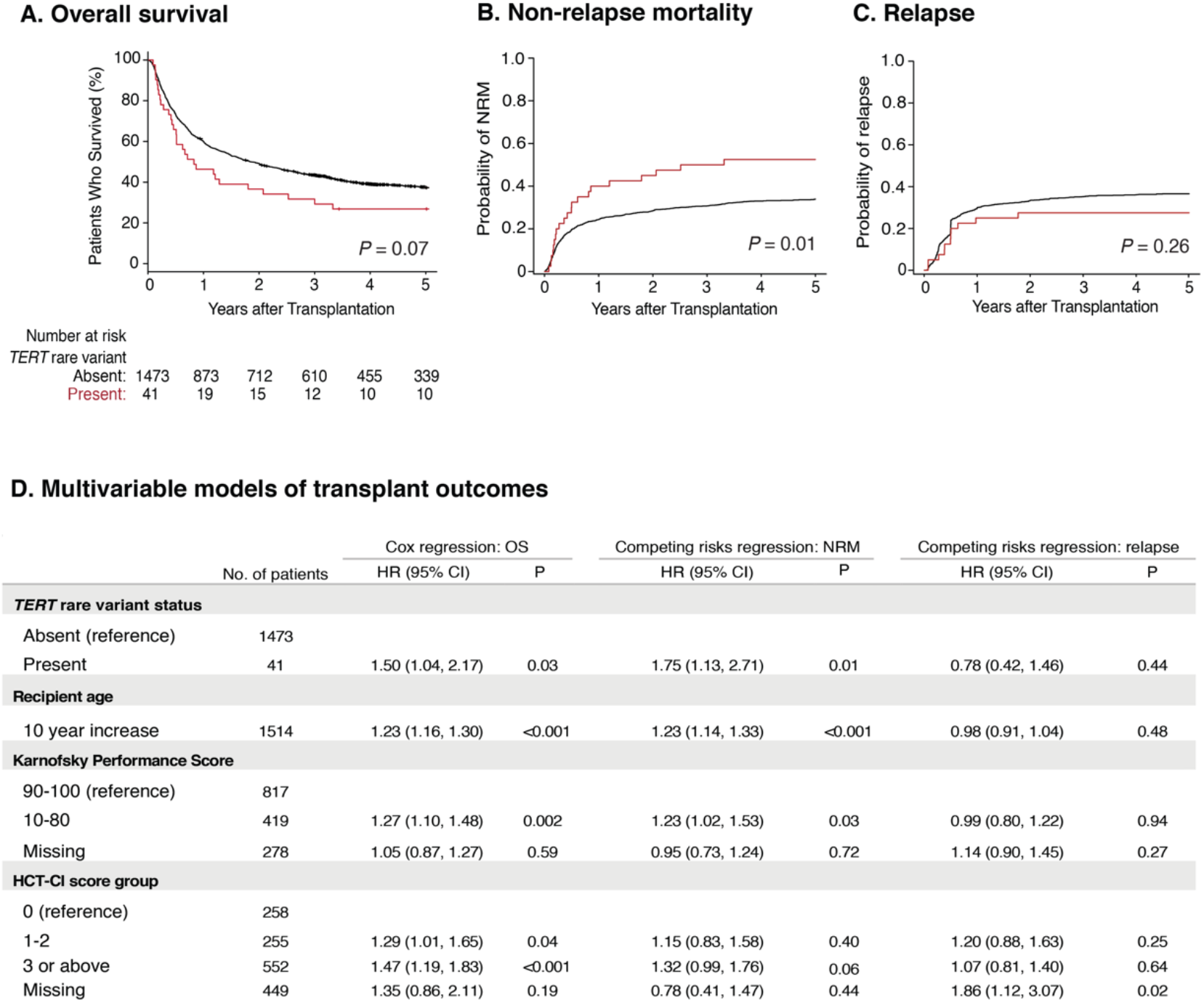
Transplant outcomes by *TERT* rare variant status. A) Kaplan-Meier curve for overall survival. B) Cumulative incidence curves for non-relapse mortality. C) Cumulative incidence curves for relapse. *TERT* rare variants are colored in red and patients without a *TERT* rare variant are colored in black. D. Multivariable models of overall survival (left), NRM (middle), and relapse (right).

Primary disease (32%), non-infectious pulmonary causes (21%), and infections (18%) were the most common causes of death in patients with a *TERT* rare variant (Table S7). Among these, non-infectious pulmonary causes of death occurred more frequently in patients with a *TERT* rare variant compared to those without a *TERT* rare variant. Five of six patients with a *TERT* rare variant and non-infectious pulmonary cause of death received myeloablative conditioning. The incidence of pre-transplant pulmonary dysfunction, as defined by HCT-CI criteria^52^, was not significantly different between *TERT* rare variant groups (p=0.27).

### Functional effects of TERT rare variants

We next determined the impact of *TERT* rare variants on telomere elongation in human cells (Figure 4A). Doxycycline-inducible expression of wild type TERT in K562 AML cells resulted in progressive increase in telomere length over 27 days, while expression of luciferase or a known catalytically impaired *TERT* variant (TERT^V694M^)^15,40^ resulted in minimal change in telomere length (Figure 4B). Bulk cell lines were generated for all 39 *TERT* missense rare variants and one *TERT* common variant (p.A279T). TERT expression was consistent throughout the experiment (Figure 4A and S6). Telomere elongation capacity was calculated as the change in qPCR telomere length from day 0 to day 27 normalized to that of wild type TERT (Figure 4C and S7).

**Figure 4.**
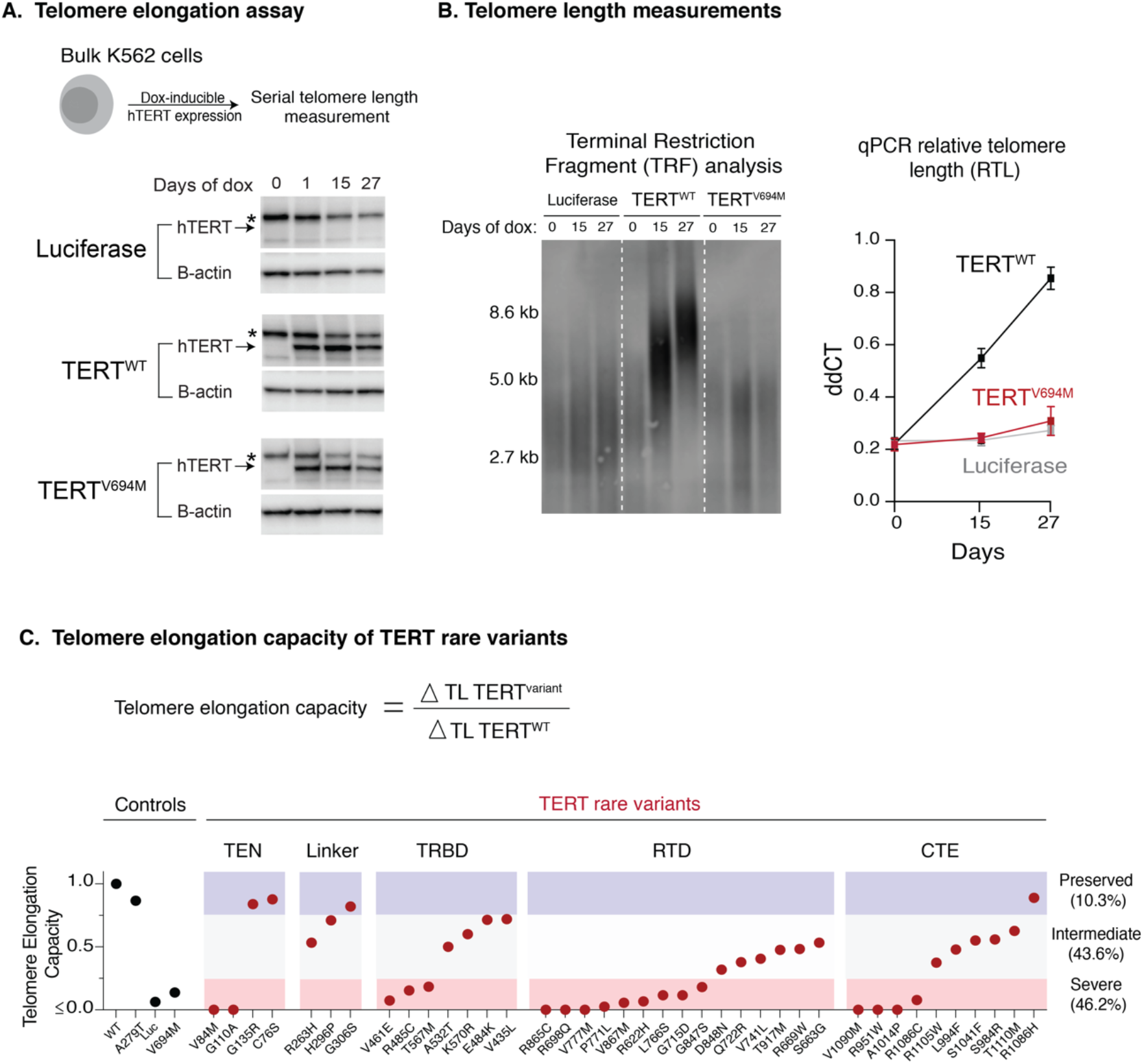
Functional characterization of *TERT* rare variants. A) Cell-based telomere elongation assay in isogenic bulk K562 cell lines with doxycycline-inducible TERT expression. hTERT expression throughout the experiment is shown for control conditions: luciferase, TERT^WT^, TERT^V694M^. hTERT band (~127kDa) is labelled with an arrow and the asterisk corresponds to a non-specific band seen in all conditions. B) Telomere length measurements by terminal restriction fragment analysis and qPCR for luciferase (gray), TERT^WT^ (black) TERT^V694M^ (red). C) Telomere elongation capacity of *TERT* rare variants normalized to wild-type TERT rare shown in ranked order grouped by structural domain. Control conditions are colored in black and *TERT* rare variants in red.

Most TERT rare variants exhibited impaired capacity to elongate telomeres compared with wild type TERT (Figure 4C; Figures S3 and S5). Eighteen variants (46.2%) displayed severely impaired telomere elongation capacity (<25% of wild type), including 10 of 11 previously reported variants associated with telomere biology disorders (Table S2). Intermediate telomere elongation capacity (25-75% of wild type) was observed for seventeen variants (43.6%). C76S, G135R, G306S, and R1086H exhibited preserved telomere elongation capacity (>75% of wild type). Severely impaired variants occurred within all major structural domains of TERT: TEN (2 of 4), TRBD (3 of 8), RTD (9 of 15), CTE (4 of 10). As a group, linker variants had a modest impact on telomere elongation capacity (H263H: 53%, H296P: 71%, G306S: 82%).

### Structural analysis of TERT rare variants

To study the potential structural effects of *TERT* rare variants, we constructed a homology model of human telomerase bound to the OB domain of the shelterin protein TPP1 as described in methods and supplement. In this model, the TRBD, RTD, and CTE domains form a closed ring structure that is consistent with low-resolution cryo-EM data from human telomerase (Figure 5A).^53^ The TEN domain straddles the insertion in fingers domain (IFD) portion of the RTD and contacts the CTE domain, thereby trapping the TERC template and DNA substrate within the active site (TERC and DNA omitted in Figure 5A for simplicity). The TPP1 OB domain docks at the TEN-IFD interface of TERT, as described previously.^53^

**Figure 5.**
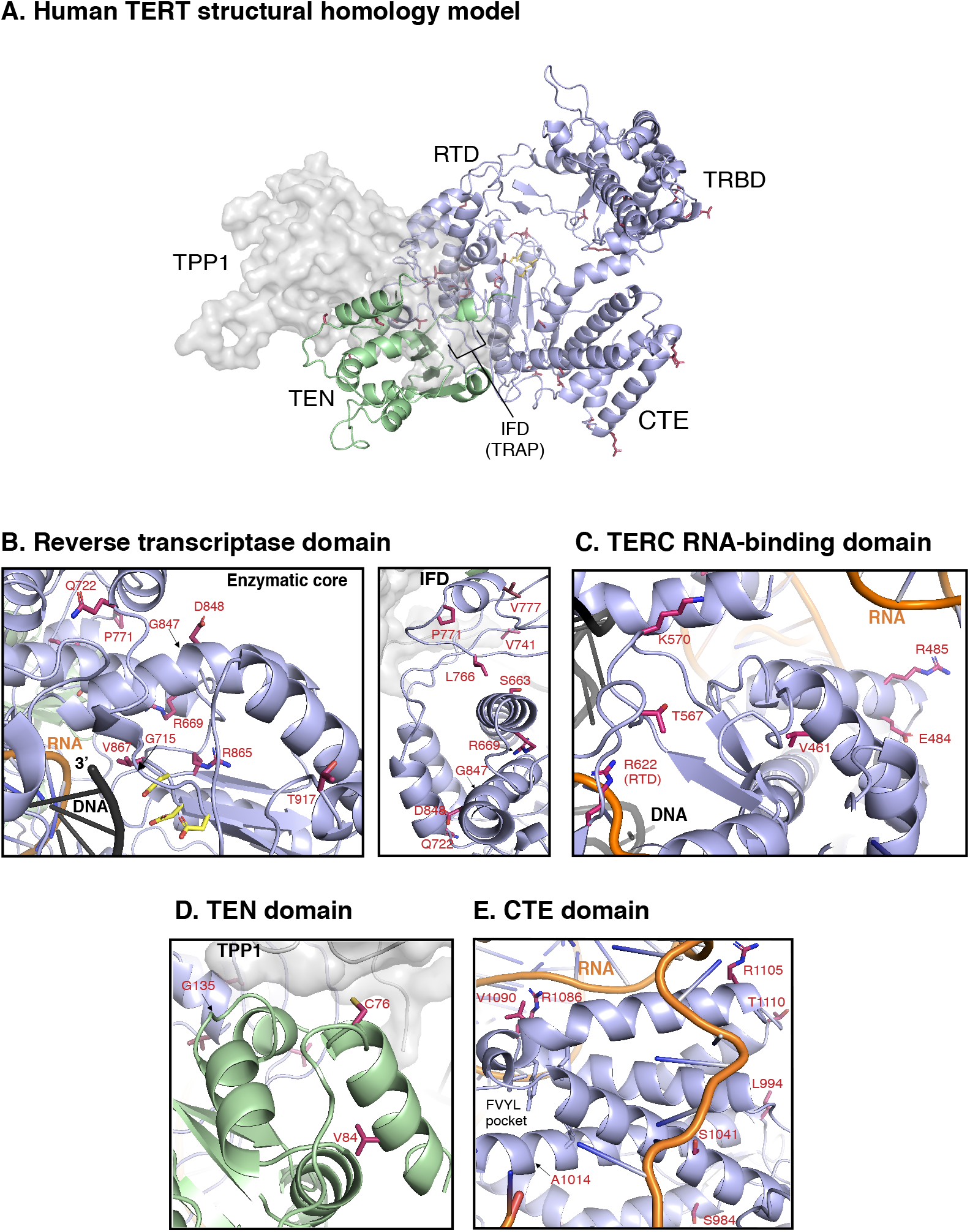
Structural analysis of *TERT* rare variants. A) Human TERT homology model. The ring, formed by the TRBD, RTD, and CTE domain is colored in purple and the TEN domain in green. Panels B, C, D, and E show the *TERT* rare variants within the RTD (enzymatic core and IFD regions), TRBD, TEN, and the CTE domains (including the FVYL pocket), respectively. Note that R622 belongs to the RTD but is displayed in panel C due to its proximity to the TRBD. The side-chains for the residues mutated in the MDS rare variants (carbon atoms colored crimson) and the catalytic residues in the active site (carbon atoms colored yellow) are shown in stick representation. The modeled regions of TERC are in orange while the DNA substrate is depicted in black.

Of the fifteen *TERT* rare variants that map to the RTD, ten localize within the catalytic core region (Figure 5B) that is structurally similar to other known reverse transcriptases. ^31^ Four of these ten variants (R622H, G715D, R865C, and V867M) are proximal to the active site pocket and the RNA-DNA duplex (*left* panel of Figure 5B; see Figure 5C for R662). In contrast, variants S663G, R669W, R698Q (not shown), G847S, D848N, and T917M map distal to the active site (*center* panel of Figure 5B; see *left* panel for T917). We also identified five RTD variants that lie within the IFD.^53^ The IFD consists of two bracing helices with an intervening TERT-specific “TRAP” subdomain that entraps the template-primer duplex within the active site (center panel of Figure 5B). Q722R resides at the base of the N-terminal bracing helix, and four variants (V777M, P771L, V741L and L766S) lie within the TRAP region. Notably, V777 resides within an α helix that contacts the TEN domain, while P771 lies immediately adjacent to it.

The TEN domain facilitates telomere repeat addition processivity (RAP) of telomerase and mediates telomerase recruitment to the ends of chromosomes through specific interactions with the TERT IFD-TRAP and the N-terminal OB domain of TPP1.^53^ Four variants (C76S, V84M, G110A (not modeled), and G135R) localize to the TEN domain and are proximal to a region implicated in recognizing TPP1 (Figure 5D).

The CTE domain, known as the thumb domain in other polymerases, is composed of four highly conserved motifs (E-I, E-II, E-III, E-IV) essential for biological activity through RNA-DNA duplex binding as well as RAP.^54^ Eight of ten CTE variants localize to E-I (R951W, S984R, L994F, A1014P), E-II (S1041F), and E-III (R1086H, R1086C, V1090M) (Figure 5C). At opposite ends of a loop connecting E-III and E-IV, R1105W sits in close proximity to TERC, whereas T1110M resides on a solvent-exposed face.

The TRBD interacts extensively with TERC via the CR4/5 domain and pseudoknot/template region.^46^ Among the eight TRBD variants, R485C, E484K, and A532T (not shown) localize to alpha-helices in contact with the CR4/5 domain of TERC in the homology model, whereas T567M and K570R reside on a hydrophilic loop in close proximity to the RNA-DNA duplex (Figure 5C). V461E targets a residue buried within the protein hydrophobic core. In contrast, variant V435L (not shown) occurs at a poorly conserved residue within an unstructured region. The remaining three variants (R263H, H296P, and G306S) localize to a weakly conserved linker region that connects the TEN domain to the TRBD (not modeled).

## DISCUSSION

The prevalence, prognostic significance, and functional effects of *TERT* variants in MDS patients have not been systematically evaluated. In this study, we identified all *TERT* variants in a registry-level cohort of 1514 MDS patients unselected for suspicion of telomere biology disorder and studied their clinical and functional consequences. *TERT* variants that are frequent in population databases, such as A279T, H412Y, E441del, and A1062T, had no apparent phenotypic consequences, consistent with studies showing that common polymorphisms do not contribute measurably to telomere-related diseases.^40^ In contrast, *TERT* rare variants (<0.1% in all reference populations) were present in 2.7% of MDS patients and were associated with characteristics of disease-causing germline mutations, including shorter telomere length, younger age at MDS diagnosis, and impaired telomere elongation capacity in human cells.

As a group, patients with a *TERT* rare variant had poor survival after allogeneic HSCT owing to an increased risk of NRM. The prognostic impact of *TERT* rare variants was independent of established clinical predictors of NRM, such as recipient age, Karnofsky performance status, HCT-CI score, donor-recipient HLA matching, and conditioning intensity. In particular, patients with a *TERT* rare variant were more likely to die from a non-infectious pulmonary cause than those without a *TERT* rare variant. This increased risk of NRM and post-transplantation pulmonary complications evokes studies that have reported high rates of NRM and fatal post-transplant pulmonary complications in patients with dyskeratosis congenita.^2,55–58^ In this context, a global defect in telomere maintenance and constitutionally short telomeres may render patients susceptible to non-hematopoietic end-organ toxicity after conditioning with radiation or DNA alkylating agents.

Most germline *TERT* variants observed in telomere biology disorder patients are missense substitutions.^32^ In the absence of compelling family history or functional data, novel variants thus present a clinical dilemma, where accurate variant classification relies on multi-tiered evidence to support interpretation.^33,34,59^ Most of the *TERT* rare variants we identified in this MDS cohort were classified as VUS and would not be definitively actionable in clinical practice. *In silico* modeling approaches have shown limited utility in predicting the effects of missense variants in patients.^35^ Moreover, *in vitro* functional assays may fail to reveal non-enzymatic defects that are essential for *in vivo* activity, including nuclear localization,^60^ holoenzyme assembly,^61^ and telomerase recruitment to telomeres.^53,62^ We therefore determined the functional effects of all candidate *TERT* rare variants in a cell-based assay with *in vivo* telomere extension as the read-out. Using this approach, we showed that 90% of rare variants caused a quantifiable defect in telomere elongation compared to wild type TERT, whereas the SNP A279T had preserved function. The degree of impairment among rare missense substitutions was variable, with 18 having severe functional effect (<25% of wild type telomere elongation capacity) and 17 having intermediate function (25-75% of wild type). Variants that displayed preserved telomere elongation capacity in our assay (>75% of wild-type) may represent private genetic variants with no significant biological impact. Conversely, a mild functional impairment of these variants may not be evident in our assay, but nevertheless contribute to clinical disease due to genetic anticipation or in combination with other factors increasing hematopoietic cell turnover.

*TERT* rare variants were distributed across multiple domains, suggesting that there are multiple mechanisms by which TERT variants can impair telomere extension. By correlating the functional data with the human telomerase homology model, we inferred the mechanistic basis of each variant’s effect. For example, while all 15 *TERT* rare variants within the RTD demonstrated reduced telomere extension, 10 were in proximity to the active site and potentially impair catalysis directly,^15^ while the 5 IFD variants likely alter TPP1-dependent telomere association.^63^ In this regard, the two variants within the TEN domain that showed severely reduced telomere elongation capacity (V84M and G110A) are also positioned to disrupt TPP1-mediated telomerase recruitment.^53^ In contrast, variants within the TRBD and CTE likely impair interactions with regions of TERC, including its CR4/5 and template/pseudoknot domains, that are important for both ribonucleoprotein complex stability and catalysis.^30^ Our complementary functional and structural analysis provides a powerful framework for evaluation of novel *TERT* variants and may enhance establishment of TERT-specific variant classification guidelines with broad clinical applicability.

Together, our results indicate that *TERT* rare variants identify a group of MDS patients who may have an unrecognized telomere biology disorder. This conclusion is based on functional characterization of all candidate variants, telomere length measurements in primary patient samples, and annotation of clinical characteristics including age, comorbidities, and toxicity outcomes in a registry-level cohort. Our analysis is limited by the unavailability of germline reference tissue and absence of detailed family history and clinical examination. Importantly, no patient with a *TERT* rare variant had a clinical diagnosis of dyskeratosis congenita, and pre-transplant clinical characteristics such as pulmonary and hepatic function, peripheral blood counts, and history of aplastic anemia were similar among patients with or without a *TERT* rare variant. This observation is consistent with previous reports that adults with telomere biology disorder rarely exhibit syndromic features and affected patients often lack a family history of hematologic, pulmonary, or hepatic abnormalities.^4,21^ Indeed, MDS has been reported to be a late presenting disease manifestation in patients with telomere biology disorders.^2,4,21^

Our results are consistent with the varied clinical presentation of patients with a telomere biology disorder and indicate that clinical criteria alone may be inadequate to identify all patients with germline *TERT* mutations. Notably, 90% of MDS patients with *TERT* rare variants were adults older than 40 years of age and there appeared to be no upper age limit. Further, the predictive value of telomere length thresholds in identifying patients with a *TERT* mutation has been shown to be poor in older patients, where the telomere length of affected patients overlaps with the lower range of the normal aging control population.^21^ The unexpectedly high prevalence of unrecognized and clinically significant telomerase alterations among adult MDS patients thus raises the possibility that routine *TERT* sequencing should be incorporated into standard MDS diagnostic evaluation irrespective of age, clinical presentation, or family history. The results of screening could directly inform transplant donor selection by enabling exclusion of candidate related donors who share the germline allele. Further, pre-transplantation referral for evaluation of comorbid pulmonary or hepatic disease and mandating less intensive conditioning regimens could mitigate the elevated risk of NRM. Such a strategy would require multidisciplinary assessment and gene-specific guidelines for variant classification to guide clinical decision-making.^64^

The frequency of *TERT* rare variants does not fully account for the adverse effect of short telomere length on non-relapse mortality in adult MDS patients.^8^ Genetic alterations to other components of the telomerase and shelterin complexes are also associated short telomeres and clinical disease.^1,2,32^ Unbiased sequencing of these genes, paired with telomere length measurement and clinical outcomes, may reveal additional gene variants associated with MDS predisposition and similarly inferior transplant outcomes.

In conclusion, we show that *TERT* rare variants impair telomere elongation in cells and are associated with shorter telomeres, younger age at diagnosis, and an increased risk of NRM in MDS patients undergoing allogeneic transplantation. These results suggest that unrecognized telomere biology disorders contribute to the pathogenesis of MDS. Identifying *TERT* variants via systematic genetic screening in MDS patients of all ages and regardless of clinical suspicion could impact clinical care by informing donor selection, family counseling, and mitigation of NRM risk.

## Methods

### Patients and samples

We previously described a cohort of 1514 MDS patients who were enrolled in The Center for International Blood and Marrow Transplant Research repository and research database who had banked whole peripheral blood DNA samples.^8^ The median follow-up time for censored patients was 5.0 years. A separate cohort of 401 adult patients with non-Hodgkin lymphoma (NHL) patients treated with high-dose chemotherapy with autologous stem cell rescue at the Dana–Farber Cancer Institute had banked mobilized whole peripheral blood DNA samples.^41^ This study was conducted with the approval of the institutional review board at the Dana–Farber Cancer Institute.

### DNA library preparation, sequencing, and variant annotation

In the MDS cohort, we sequenced the *TERT* coding region (exons 1-16) and known germline single nucleotide polymorphisms (SNPs). In the NHL cohort, we sequenced *TERT* in DNA extracted from mobilized whole peripheral blood samples. The genetic analysis was completed and locked prior to merging with clinical data. Detailed sequencing methods are in the supplementary methods.

### Telomere Length Measurements

Relative telomere length of MDS patients was measured by quantitative polymerase chain reaction (qPCR) as previously reported.^8^ K562 cell DNA was extracted using the QIAamp Blood Mini kit (Qiagen). Telomere length was measured by two orthogonal methods: 1) qPCR^8^ and 2) telomere restriction fragment (TRF) analysis using TeloTAGGG Telomere Length Assay (Sigma Aldrich) according to the manufacturer’s protocol.

### Plasmids, cloning, site-directed mutagenesis

Human TERT cDNA^39^ was cloned into the gateway pDONR221 plasmid (Invitrogen). *TERT* variants were generated using the Q5 Site-Directed Mutagenesis Kit (NEB) and primers designed using NEBase changer (Table S1). Transfer of each construct to the pCW57.1 destination plasmid (Addgene) was performed using LR clonase (ThermoFisher) and the complete sequence was confirmed by Sanger sequencing.

### Lentivirus production and cell line generation

Lentivirus for each *TERT* variant was produced in HEK 293T cells. *TP53*-repaired K562 cells (gift from the Ebert lab)^42^ were transduced at MOI ~ 1 followed by puromycin selection (2ug/mL) to generate bulk cell lines. K562 cells were treated with doxycycline (1ug/mL) for 27 days. RPMI media was added every 2 days and cells were split every 3 days.

### Western blotting for hTERT expression

Protein extracts were prepared in Laemmli 2X buffer and western blots performed using SDS gels (BioRad) and Trans-Blot Turbo Transfer System. hTERT expression was visualized using hTERT antibody (Rockland; Cat# 600-401-252; 1:1,000 dilution) with anti-rabbit secondary antibody. B-actin was used as loading control (Abcam; Cat# ab20272; 1:10,000 dilution). Chemiluminescence images were obtained using Bio-Rad ChemiDoc and analyzed using Image Lab software.

### Structural modeling of residues mutated in TERT rare variants

A homology model of human telomerase bound to TPP1-OB and parts of TERC was generated using the cryo-EM structure of the *Tetrahymena thermophilia* telomerase holoezyme^43,44^, the crystal structure of human TPP1-OB^45^ (as described in Tesmer and Smith et al, *PNAS* 2019), as well as additional crystal structures of TERT/TERC domains from various species^43,45–47^ using Phyre 2.^48^ *TERT* rare variant positions were manually annotated in the final homology model. Details of the homology model are listed in supplemental methods.

### Statistical analyses

Fisher’s exact test was used to test for association between pairs of categorical variables. The Wilcoxon rank-sum test was used to assess a location shift in the distribution of continuous variables between two groups. For associations with ordered categorical variables, Cochran-Armitage trend test was used for singly-ordered contingency tables. All p-values were two-sided.

Overall survival was defined as the time from transplant until death from any cause. Subjects not confirmed dead were censored at the time last known to be alive. Differences in survival curves were assessed using log-rank tests. NRM was defined as death without relapse. NRM, with relapse as a competing risk, was assessed with the use of Gray’s test. For relapse, death without relapse was considered a competing risk. Univariate and multivariable analyses of OS were performed using Cox regression. OS estimates were calculated using the method of Kaplan-Meier and reported with 95% CIs based on Greenwood’s formula. Hazard ratios with 95% CIs and Wald P values were reported for covariates in multivariable Cox models. Multivariable models for competing risks of relapse and NRM were generated using the Fine and Gray method.

## Acknowledgements

This work was supported by the National Institutes of Health grants K08CA204734 (R.C.L.), RC2DK122533 (R.C.L.), R01AG050509 (J.N.), and R01GM120094 (J.N.), the Aplastic Anemia & MDS International Foundation (R.C.L.), the Jim and Lois Champy Fund (R.C.L.), the Anna Fuller Fund (R.C.L.), American Cancer Society Research Scholar grant RSF-17-037-01-DMC (J.N.), National Institutes of Health (NIH)/National Heart, Lung, and Blood Institute fellowship training grant 2T32HL116324-06 (C.R.R.), the Sigrid Juselius Foundation (M.M.), the Maud Kuistila Memorial Foundation (M.M.), the Väre Foundation for Pediatric Cancer Research (M.M.) the Orion Research Foundation (M.M.), T32-CA009172 (F.D.T.) the Jock and Bunny Adams Education and Research Fund (J.H.A.), NIH/National Institute of Diabetes and Digestive and Kidney Diseases 1R01DK107716 (S.A.), and the Dana-Farber/Harvard Cancer Center core grant from the National Cancer Institute 5P30 CA006516 (R.R. and D.N.). The Center for International Blood and Marrow Transplant Research is supported primarily by Public Health Service Grant/Cooperative Agreement 5U24CA076518 from the National Cancer Institute, the National Heart, Lung and Blood Institute, and the National Institute of Allergy and Infectious Diseases (W.S., S.R.S. and Z.-H.H.); grant/cooperative agreement 4U10HL069294 from the National Institutes of Health, National Heart, Lung and Blood Institute and National Cancer Institute; Health Resources and Services Administration contract HSH250201200016C; and grants N00014-18-1-2850 and N0014-19-1-2888 from the Office of Naval Research. The views expressed in this article do not reflect the official policy or position of the National Institutes of Health, the Department of the Navy, the Department of Defense, Health Resources and Services Administration or any other agency of the US Government.

## Authorship Contributions

M.M., C.R.R., and R.C.L. designed the study, analyzed data, and wrote the manuscript; C.J.G., M.M.C., E.H.O., and I.D.V. performed sequencing; D.N. and R.R. curated clinical data, conceived the statistical plan, and performed statistical analysis; C.R.R., H.Q.R, and J.E.G performed consensus variant classification. V.T., S.P., J.N. performed structural modeling, analyzed data, and contributed to research discussion; F.D.T. and D.K. performed cloning and generated cell lines; W.S., S.R.S., and Z.-H.H. curated clinical data, clinical data collection and quality assurance, and contributed to research discussion; S.A. C.C., and J.H.A. interpreted data and contributed to research discussion; and all authors reviewed the manuscript during its preparation and approved the submission.

## Disclosure of Conflicts of Interest

D.N. has received research support from Pharmacyclics and owns stock in Madrigral Pharmaceuticals. M.M. has received honoraria from Celgene and Sanofi. R.C.L. has received research support from Jazz Pharmaceuticals and consulting fees from Takeda Pharmaceuticals and bluebird bio.

## Notes

### Summary of Updates

Re-wrote the abstract. Made figures more clearly presented. No changes to main text or figure content.

